# Promoter Structural Variants are Drivers of Genome-Wide Differential Expression in Maize

**DOI:** 10.64898/2026.06.23.734061

**Authors:** Manisha Munasinghe, Andrew Read, Aimee Schulz, Yaniv Brandvain, Nathan Springer, Candice Hirsch

## Abstract

**Background:** Structural variants (SVs) are large insertions or deletions of DNA sequences. While less numerous than single nucleotide polymorphisms, SVs often account for a greater proportion of nucleotide differences between genomes. Their size and frequent association with repetitive sequences has historically hindered their detection, which has limited the ability to associate this variation with molecular and phenotypic trait variation. While some SVs have been linked to observable traits, it remains unclear whether such effects are rare or broadly distributed across the genome.

**Results:** To test for genome-wide relationships between SVs and gene expression, we analyzed genome assemblies and transcriptomic data from 10 tissues across 26 diverse maize inbred lines. We identified SVs amongst these lines and examined variants located within the 1kb promoter region upstream of genes. Thousands of genes showed expression differences associated with promoter SVs, often in a tissue-specific manner. One common feature of these SVs was the presence of transposable element sequences. LTR retrotransposons were enriched amongst promoter SVs associated with differential expression and often reduced expression of the nearby gene. Despite widespread expression changes, we found no enrichment for specific biological functions or pathways among affected genes.

**Conclusions:** Our findings indicate that extant TE-mediated promoter SVs play a significant role in shaping gene expression patterns across the maize genome. However, their phenotypic effects appear limited or context-dependent, suggesting that many variants may have minimal impact outside specific developmental stages or environmental conditions.

## Background

Recent advancements in computational genomics have enabled researchers to investigate structural variation between genomes at an unprecedented scale and resolution [1]. While the minimum size requirement may vary (>10, 20, or 50 bp)[2], structural variants (SVs) are generally defined as insertions or deletions of DNA sequences that are at least 50 base pairs in length. The size of SVs can vary widely from just over 50 base pairs to several megabases. While single nucleotide polymorphisms (SNPs) are more numerous, SVs typically account for a greater number of nucleotide differences between genomes [3–5]. Despite their importance, SVs have historically been difficult to study due to technical limitations. Short-read sequencing technologies often struggle to accurately identify SVs, largely due to their size and frequent association with repetitive sequences, both of which complicate read mapping and assembly. Long-read sequencing has addressed some of these challenges, but technical and computational difficulties still restrain their usage [6–8]. Pan-genomic approaches, which often integrate both short- and long-read data to construct multiple high-quality genome assemblies, have greatly reduced the technical barriers to studying SVs throughout the genome [9,10]. As a result, pan-genomic studies have become a powerful tool for detecting structural variation and linking them to phenotypic traits [11].

Our historic inability to reliably detect SVs has hindered a comprehensive understanding of the extent of structural variation between genomes and their contribution to phenotypic diversity. Still, across major crop species, SVs have been implicated in key agronomic traits, such as flowering time, frost tolerance, and yield-related characteristics [12–15]. In avian systems, structural variation has been associated with differences in body weight, plumage colouration, and even behavior [16]. Perhaps the most thoroughly studied examples come from humans, where SVs have been linked to nearly every category of genetic disorder – including Mendelian diseases, complex traits, and metabolic conditions [17,18]. This growing body of work clearly demonstrates the potential of SVs to influence phenotypic variation. However, it remains unclear whether such effects are widespread across the genome or if these examples represent isolated cases. There are many mechanisms by which SVs could result in a phenotypic effect. SVs within exons or coding regions can disrupt expression by knocking out the gene entirely or by altering transcript structure [19]. However, a recent multi-tissue analysis of human gene expression patterns found that causal SVs were most frequently noncoding variants located within regulatory regions [20]. While these findings are compelling, our understanding of how SVs in regulatory regions influence gene expression remains incomplete. The effects of such SVs likely depend on the extent to which they disrupt interactions between genes and their regulatory elements [21–24] and may also be influenced by the sequence content of the variant itself (e.g. SVs containing transposable elements).

Transposable elements (TEs) are mobile, repetitive DNA sequences capable of replicating and inserting themselves into new genomic locations. This insertion consequently generates structural variation. When inserted into regulatory regions, TEs can influence gene expression in multiple ways. TEs often contain transcription factor binding sites, promoters, or enhancers in order to increase their transposition rate, and these regulatory features can potentially increase the expression of nearby genes [25–29]. However, host genomes have evolved a myriad of mechanisms to suppress TE activity. This includes transcriptionally silencing them using DNA methylation, histone and chromatin modifications, and small interfering RNAs [30–33]. While these mechanisms are effective at silencing TEs, they can have unintended effects on adjacent genes. The deposition of epigenetic modifications is not always precise and nearby sequences may inadvertently be repressed leading to reduced gene expression [34,35]. Moreover, chromatin modifications are dynamic and often vary in response to external stimuli. Consequently, TE insertions may not only alter gene expression but may do so in a context-dependent manner [36–39]. Despite these possibilities, the frequency and extent of context-dependent gene expression driven by TE insertions remains poorly understood. On one hand, such variation could serve an adaptive role by enabling fine-tuning of gene expression across cell types or in response to stress [40–43]. On the other hand, most disruptions to established expression patterns are likely deleterious and subject to purifying selection [44,45]. It remains an open question how often SVs, particularly those derived from TEs, contribute to context-specific regulatory changes across the genome.

Maize is a powerful system for investigating the relationship between structural variation and gene expression, as over 70% of its genome consists of annotated TE sequences [46–48]. There is also extensive genomic resources for the species, including available genome assemblies, TE annotations, epigenetic and chromatin profiles, and extensive transcriptional data for 26 widely studied and genetically diverse inbred lines [47]. Recently, base-pair resolved pairwise whole-genome alignments were generated between the B73 reference genome and the other 25 genomes [49]. This enabled an exceptionally detailed analysis of structural variation amongst these lines. From these and related datasets, we can precisely identify the breakpoints of SVs, quantify their TE content, and assess their impact on gene expression across genotypes and tissue types.

In this study, we identified SVs located within the 1kb promoter region upstream of annotated genes for the maize reference genome B73. Genotypes sharing the same SVs across the promoter and gene body were grouped into structural haplotypes for a gene, allowing us to compare gene expression between haplotypes and to assess whether promoter SVs are associated with differential expression. We identified thousands of genes with expression differences between structural haplotypes, many of which showed tissue-specificity. In addition to mean differences in expression between structural haplotypes, differentially expressed promoter SVs often exhibited larger changes in expression variance across tissues as well. Interestingly, the only common characteristic of promoter SVs associated with differential expression was an enrichment for SVs that overlap TEs. We found an excess of LTR retrotransposons in the promoters of differentially expressed genes and, when differential expression was observed, these retrotransposon sequences were more often associated with reductions in expression.

## Results

### Identification of Promoter-SV Structural Haplotypes for Expression Analyses

Structural variants were identified from pairwise whole-genome alignments between the B73 reference genome and each of the remaining 25 diverse lines. To identify shared SVs across contrasts, variants were considered comparable if they had similar start/end positions, size, and the same TE-based category as defined in Munasinghe et al (2023a) (Figure S1) [49]. Equivalent SVs were assigned a common ID enabling consistent tracking across genotypes. We generated a structural haplotype for each gene (gene body plus the upstream 1kb promoter sequence) in every line by concatenating the IDs of overlapping SVs. On average, each B73 gene had seven unique structural haplotypes with four lines per haplotype (Figure S2A,B). The mean frequency of the major structural haplotype for each gene was roughly 50% with approximately six associated minor structural haplotypes (Figure S2C).

Genes with at least two structural haplotypes in at least three lines each that had no SVs overlapping the gene body and showed evidence of expression were retained. We refer to this subset of genes as the ‘Promoter-SV’ gene set (N = 11,737). Most genes in this set (∼68%) had only two included structural haplotypes (Figure S3A) and 80% of the retained genes included 15 or more lines (Figure S3B). Among genes with only two structural haplotypes (N= 8,005), ∼27% had relatively balanced representation with haplotypes differing by five or fewer lines (Figure S3C). Additionally, we confirmed that expression levels of Promoter-SV genes were comparable to all other B73 genes, indicating that these genes are not biased towards low expression (Figure S3D).

### Genes with Promoter Structural Variants are Enriched for Differential Expression

Transcriptomic data from ten tissues across all the lines was used to test whether promoter structural variants influence gene expression using a mixed-effect linear model. Because each gene includes a unique set of structural haplotypes and lines, we assessed significance using a permutation-based approach instead of a traditional correction for multiple tests. For the Promoter-SV gene set, 3,584 differentially expressed genes were identified, which amounted to approximately 30% of the gene set and 9% of all B73 genes. To test if the 1kb size for the promoter was restrictive, the analysis was repeated with the promoter sequence extended to the 2kb region upstream of the gene body. Similar results were observed for both the 1kb and 2kb defined promoter regions (Figure S4).

To evaluate whether the level of differential expression observed exceeds what would be expected by chance, a random mimic of the data set was constructed. For each of the Promoter-SV genes, a non-tested B73 gene was randomly selected and assigned the same haplotype structure and line composition as one of the originally tested genes, but the mimicked gene retained its original expression values. We refer to this gene set as the Promoter-Random gene set. Using the same statistical test for differential expression, only 639 differentially expressed genes were identified, an approximately 5.5-fold reduction in comparison with the Promoter-SV gene set. This indicates that promoter SVs are associated with differential expression more frequently than expected by random chance (Figure 1A).

**Figure 1.**
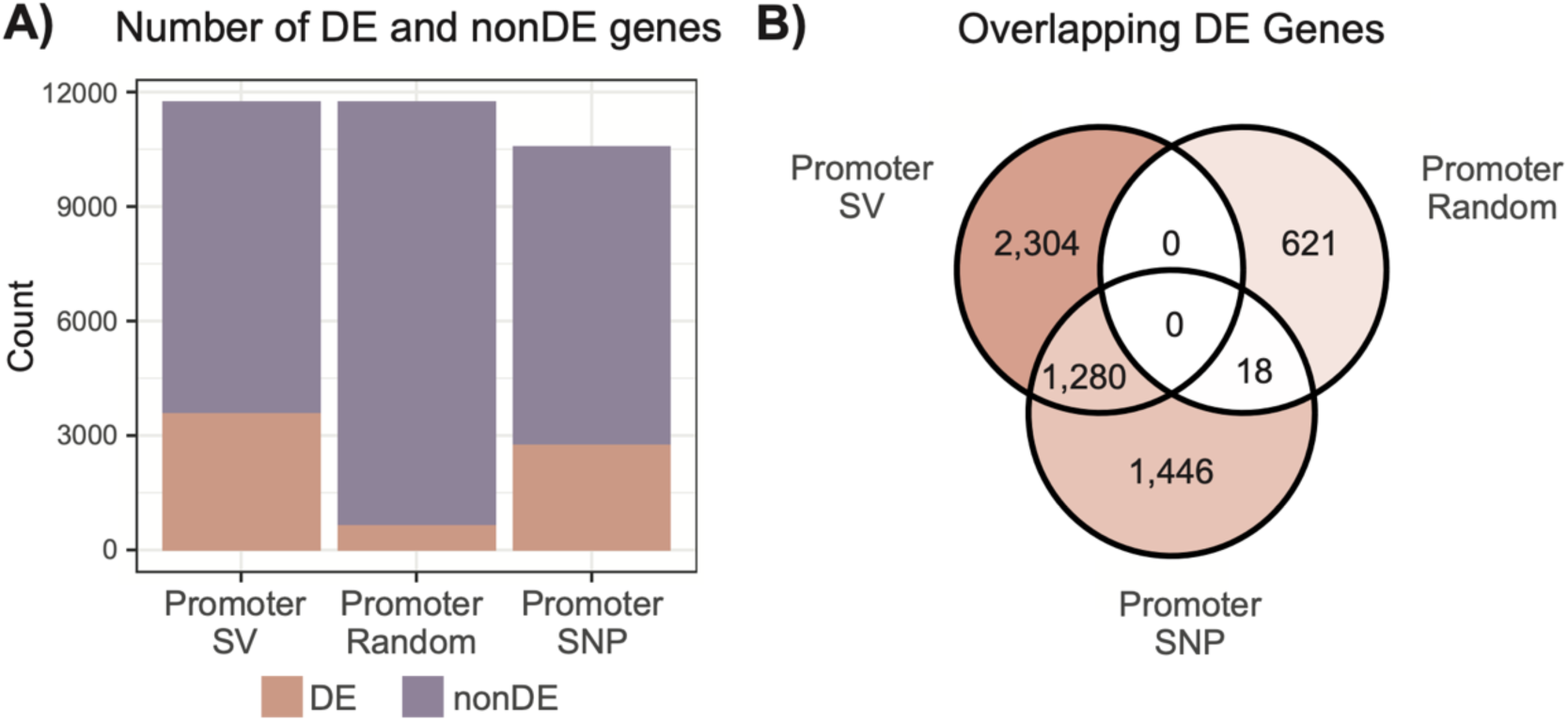
Promoter Structural Variation is Enriched for Associations with Differential Expression. (A) Barplots showing the number of differentially expressed (DE) and non-differentially expressed (nonDE) genes for each tested gene set. (B) A Venn diagram showing the overlap of genes associated with differential expression between tested gene sets.

It is important to note that in some cases, observed expression differences may actually be driven by other variants that happen to co-segregate with the structural haplotypes being tested. These other variants could be SNPs or small InDels within the promoter or coding region, as well as those that may be further upstream of the gene. In such cases, the structural haplotype will be statistically correlated with expression variation but not causally responsible for it. This possibility cannot be ruled out on a gene-by-gene basis, but it is unlikely to be a pervasive issue. Therefore, it is appropriate to examine whether general patterns emerge amongst promoter SVs that are associated with differential expression.

To test whether the observed expression differences could instead by explained by SNP variation within the promoter instead of SVs, SNP haplotypes were created using SNP variation within the 1kb promoter instead of SV content. All other criteria remained the same (i.e., no SVs within the gene body and minimum expression requirements). We refer to this gene set as the Promoter-SNP gene set. This yielded 10,576 genes, including 5,855 shared with the Promoter-SV gene set. After applying the same model with SNP haplotypes in lieu of structural haplotypes, 2,744 differentially expressed genes were identified. This is approximately 26% of the Promoter-SNP gene set, which is slightly smaller than the proportion of significant genes for the Promoter-SV gene set. Of these, 1,280 of these genes were significant in both the Promoter-SV and the Promoter-SNP gene sets (Figure 1B).

Of the 1,280 overlapping genes, only 37 genes shared identical haplotype structures between the SNP and SV analyses. On average there are more SNPs than SVs within the promoter causing the haplotypes to be split apart more with SNPs than with SVs. As such, all the lines for a SNP haplotype may perfectly overlap a structural haplotype but not vice versa. Indeed, we did find 3,676 genes that met this criteria, 1,299 of which were differentially expressed between structural haplotypes (∼36% of the identified differentially expressed genes). This suggests that while the gene-level overlap in significance is relatively high, the haplotype compositions used in each analysis are often distinct or only partially overlapping. Taken together, these results support the conclusion that promoter structural variation is associated with differential gene expression more frequently than expected by chance and that these differences are unlikely to be completely driven by co-segregating SNP variation in the same region.

### Differential Expression Exhibits Tissue-Specificity

The previous analysis only permuted the haplotype, and, as such, significance for the effect of tissue could not be determined. To explore tissue specificity, we took a tissue-by-tissue approach where we tested whether there was differential expression between structural haplotypes for each tissue. Promoter-SV genes were then grouped into three categories based on these results across all 10 tissues – Always Significant, Never Significant, or Partially Significant. Always Significant genes were those that were significant in our full analysis and significant in every tissue-specific analysis. Similarly, Never Significant genes were nonsignificant in the full analysis and in every tissue-specific analysis. Partially Significant genes were those that were significant in some but not all analyses.

Of our 11,737 Promoter-SV genes, we found that only 150 genes (∼1.3%) were Always Significant and 3,219 genes (∼27.4%) were Never Significant. The vast majority of genes (N=8,368, ∼71.3%) exhibited variable significance across the 10 tissues (Figure 2A). Of the Partially Significant genes, 3,434 were identified as differentially expressed in our full ten-tissue analysis; however, the differential expression was clearly being driven by a subset of the included tissues. We were curious whether our Partially Significant differentially expressed genes were simply losing differential expression in a handful of tissues, perhaps due to a lack of expression. However, the majority of the differentially expressed genes exhibited tissue-specific expression in only 1 – 5 tissues (Figure 2B).

**Figure 2.**
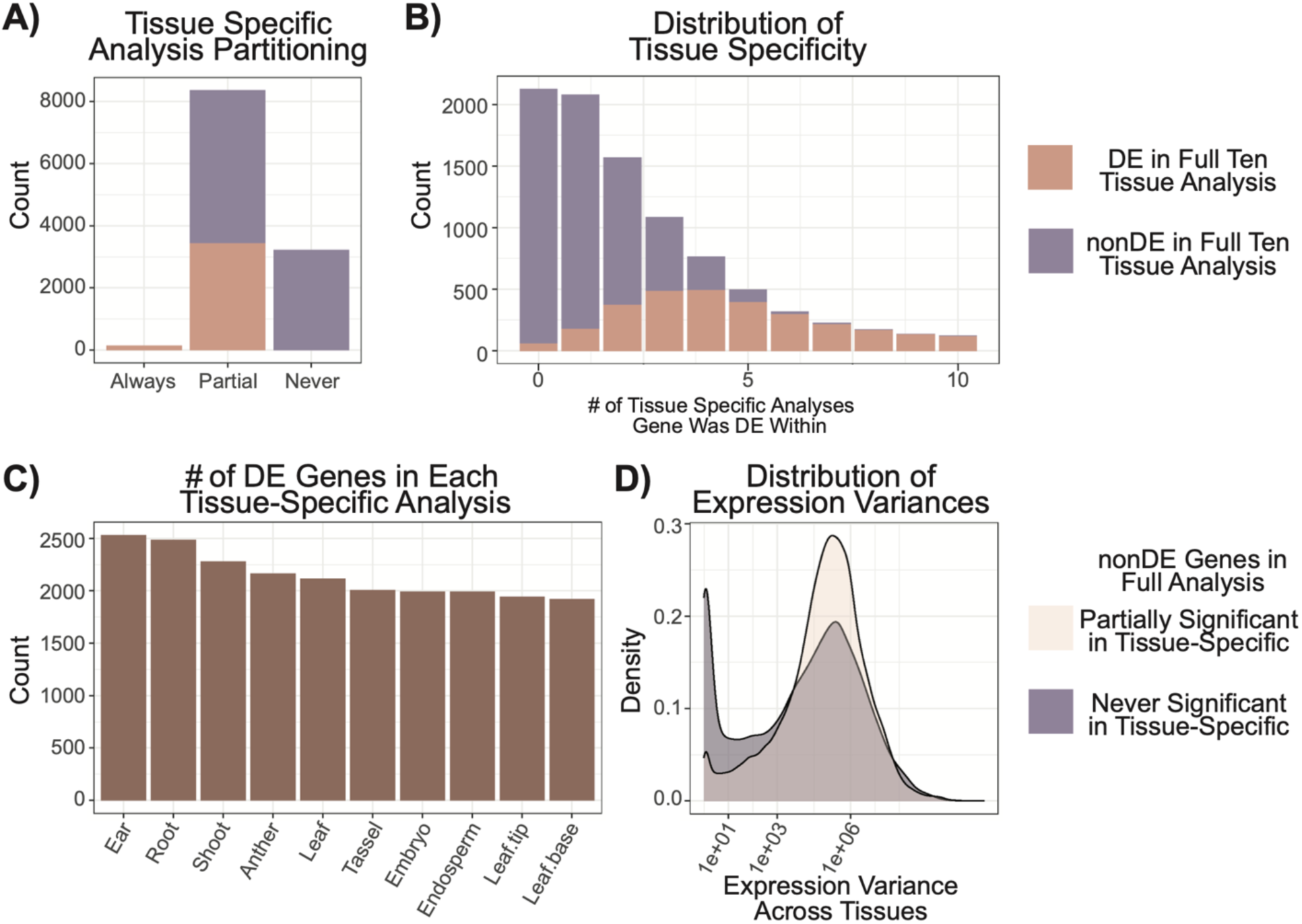
Tissue Specific Analysis of Differential Expression. (A) All tested genes are categorized as either Always Significant (differentially expressed in full ten tissue analysis and all tissue-specific analyses), Partially Significant (differentially expressed across some analyses), or Never Significant (not differentially expressed full ten tissue analysis or any of the tissue-specific analyses). (B) A distribution showing how many of the tissue-specific analyses genes were differentially expressed in. Both (A) and (B) are colored by their differential expression status in the full ten-tissue analysis. (C) A distribution showing the number of differentially expressed genes in each tissue-specific analysis. (D) For non-differentially expressed genes in the full analysis, the distribution of expression variances is colored by whether it was considered Partially Significant or Never Significant.

Given the relatively low number of significant tissues per gene, we wanted to explore if certain tissues exhibited an excess of differentially expressed genes. Due to either missing data or a lack of expression, certain genes in the full Promoter-SV gene set were not included for a given tissue. Consequently, we focus this analysis on the 9,111 genes in the Promoter-SV gene set that were tested in every tissue-specific analysis. The number of differentially expressed genes was fairly similar across the ten tissues (min= 1,920 DE genes in leaf.base, max= 2,530 DE genes in ear). The top four tissues with differentially expressed genes were ear, root, shoot and anther (Figure 2C).

Nonsignificant genes from the overall analysis that were significance in a subset of tissues were a curious set of genes that we sought to explore further. We hypothesized that this could result from high expression variance across tissues resulting in a lack of significance when all tissues were included. In this logic, removing certain tissues might amplify differences between structural haplotypes leading to significance. To test this, we calculated the variance in normalized expression across all ten tissues for each gene and compared the variance distributions between Never Significant and Partially Significant genes that were not differentially expressed in the full analysis. A statistically significant difference in the distribution of expression variance across tissues was observed (bootstrap univariate KS-test p < 1x10^-4^, Figure 2D), with Partially Significant genes showing an excess of genes with higher expression variance across tissues. This could explain why genes that were not differentially expressed in our full ten-tissue analysis exhibited significant differential expression in a specific tissue.

### Promoter SVs Contribute to Haplotype-Specific Expression Variability

Mixed-effect models evaluate signal relative to a noise. Consequently, even if mean expression differs between groups, the test statistic can become small (or nonsignificant) if the expression of a gene is highly variable within groups. We hypothesized that, instead of or in addition to shifting mean expression, promoter SVs could instead be contributing to higher expression variance akin to the SV resulting in more “chaotic” expression profiles. To explore differences in expression variance, we used two complementary variance metrics: the coefficient of variation (CoV), which captures the relative variability in expression, and the mean absolute deviation (MAD), which reflects the average absolute difference between expression values and their mean.

For each structural haplotype, the mean expression variance was calculated across all included lines. We then calculated the absolute differences in expression variances between the two structural haplotypes per gene and constructed a distribution of these absolute differences. This analysis was done on a subset of the Promoter-SV genes that had only two tested structural haplotypes differing by a single SV (N=5,872). This way differences in variance could be attributed to presence or absence of the single SV. From the distribution, genes with exceptionally similar (bottom 2.5%) or divergent (top 2.5%) expression variability between haplotypes were identified. Differentially expressed genes were significantly enriched for presence amongst the top 2.5% of genes showing differences in the mean CoV between structural haplotypes (Table 1). Conversely, differentially expressed genes were also significantly de-enriched for presence amongst the bottom 2.5% of genes showing minimal difference in the mean MAD between structural haplotypes. This suggests that differentially expressed genes more often exhibit greater variability in expression and are also more likely to have larger differences in expression relative to the mean, and that promoter SVs that result in differential expression are also more likely to increase the relative variance in expression across tissues.

**Table 1.** Chi-Square Tests for Enrichment of Differentially Expressed Genes in Extremes of Expression Variability Distributions.

| Gene List | Full Set<br>vs<br>Differentially Expressed | Full Set<br>vs<br>Non-differentially Expressed | Differentially Expressed<br>vs<br>Non-differentially Expressed |
| --- | --- | --- | --- |
| **Chi-Square**<br>**CoV p-value** | $2.90 \times 10^{-2}$ * | $3.22 \times 10^{-1}$ | $1.15 \times 10^{-3}$ ** |
| % in bottom<br>quantile | 2.5% vs 2.4% | 2.5% vs 2.5% | 2.4% vs 2.5% |
| % in top<br>quantile | 2.5% vs 3.7% | 2.5% vs 2.0% | 3.7% vs 2.0% |
| **Chi-Square**<br>**MAD p-value** | $8.85 \times 10^{-7}$ *** | $6.44 \times 10^{-1}$ | $2.211 \times 10^{-9}$ *** |
| % in bottom<br>quantile | 2.5% vs 0.38% | 2.5% vs 3.2% | 0.38% vs 3.2% |
| % in top<br>quantile | 2.5% vs 2.5% | 2.5% vs 2.5% | 2.5% vs 2.5% |

Pearson's Chi-squared test with Yates's continuity correction. \*, \*\*, and \*\*\* refer to a p-value less than 0.05, 0.01, and 0.001 respectively

Genes in the top 2.5% of expression variance demonstrated a much broader distribution of expression values with a large peak near 0 and a broad tail, while genes in the bottom 2.5% demonstrated a much more bimodal distribution with a peak near 0 and another peak near approximately 1,000 normalized counts (Figure S5). We wondered whether this pattern might reflect haplotype-specific silencing where one haplotype is transcriptionally inactive. To test this, the mean expression for each structural haplotype was compared and classified as off if the mean expression was below two normalized counts. Among genes with similar expression variance, we found 60.8% of genes were ‘on’ in both haplotypes, 37.5% were off in both haplotypes, and only 1.7% were ‘on’ in one haplotype and ‘off’ in the other. In contrast, among genes with divergent expression variance, 63.3% were ‘on’ in both haplotypes, 26.5% were ‘off’ in both haplotypes, and 10.2% were ‘on’ in one haplotype and ‘off’ in the other. This is an over 5-fold increase in the number of genes showing an ‘on/off’ difference between structural haplotypes. While this is only a fraction of all Promoter-SV genes, this pattern clearly demonstrates that promoter SVs more often result in the transcriptional silencing of one haplotype relative to the other.

### Promoter SVs Associated With Differential Expression Share Few Common Features

Shared characteristics amongst promoter structural variants that are associated with differential gene expression could suggest that SVs influence expression in predictable, repeatable ways. Conversely, a lack of common features may imply that SVs alter expression more randomly. To test this, we first compared the size distribution of promoter SVs as we hypothesized that larger SVs are more likely disrupt associations between genes and their cis-regulatory elements. Surprisingly, we found no difference in the size distribution of promoter SVs between significant and nonsignificant genes (Figure 3A). However, genes with high expression variability did have larger SVs relative to those with low expression variability (Figure S5A). Next, we looked at the distance between each promoter SV and the transcription start site (TSS) for the canonical transcript of the gene as SVs closer to the TSS might influence gene expression more directly. A modest shift towards shorter distances for significant genes was observed, suggesting that SVs associated with differential expression may lie slightly closer to the TSS (Figure 3B). These distributions are largely overlapping though, indicating that proximity alone is not a consistent predictor of expression differences.

**Figure 3.**
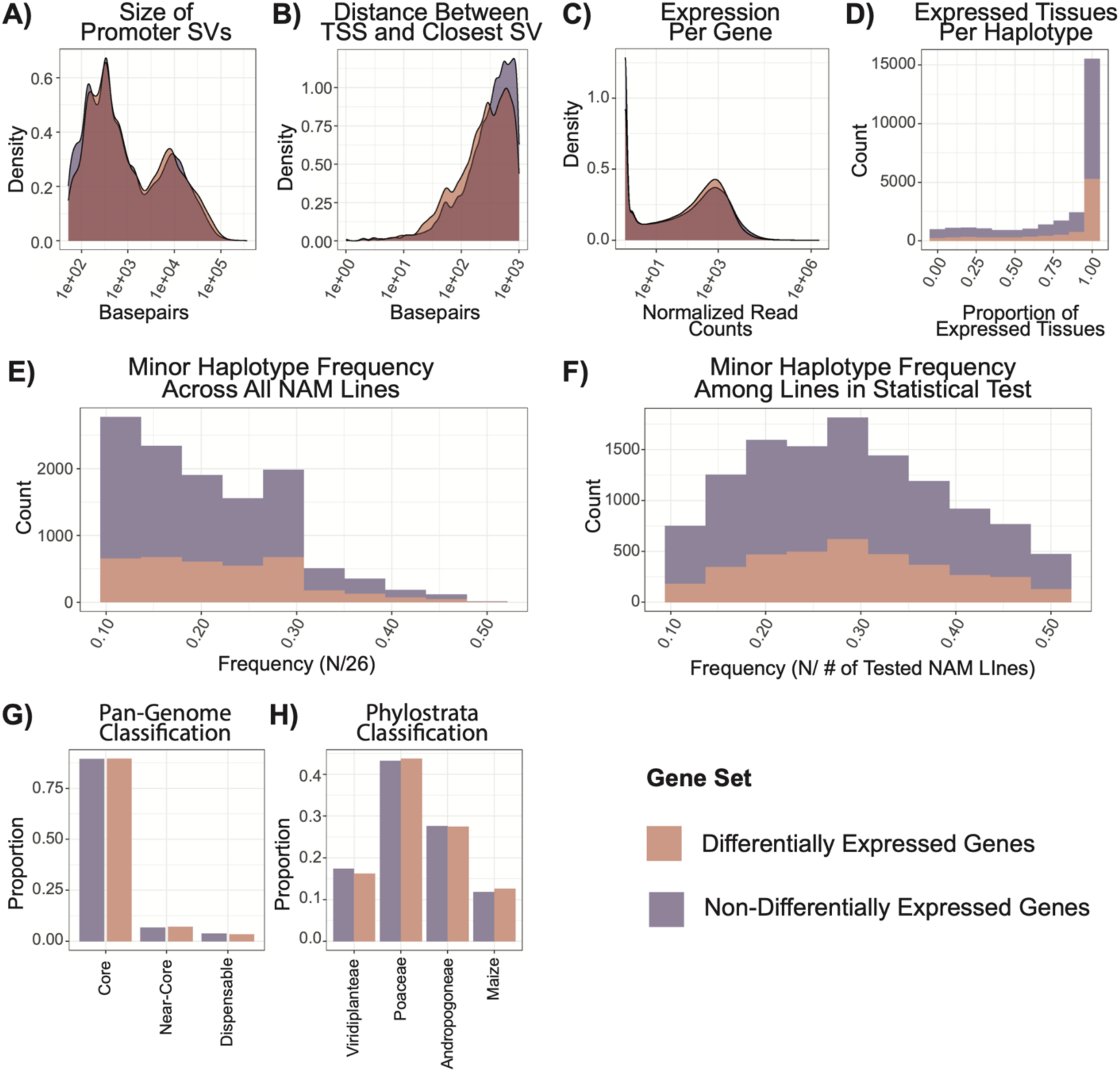
Limited Differences Between Differentially Expressed and Non-Differentially Expressed Promoter-SV Genes. Several different features were contrasted between significant and nonsignificant gene sets including (A) the size of the structural variant located within the promoter, (B) the distance between the transcription start site (TSS) and the closest promoter structural variant, (C) the distribution of gene expression values across tissues, (D) the number of expressed tissues per structural haplotype, (E) the allele frequency distribution of the minor haplotype, defined as the second most common haplotype, amongst the entirety of the lines, (F) the allele frequency distribution of the minor haplotype amongst the line including in the statistical test of differential expression, (G) the proportion of each gene set belonging to distinct pan-genome classifications including core (gene is present amongst all 26 lines), near-core (gene is present amongst 23 - 24 lines), or dispensable (shared in 2 - 22 lines), and (H) the proportion of each gene set belonging to different phylostrata moving from deeper within the clade (Viridiplanteae) to younger Maize-specific genes.

In addition to features of the promoter, we also tested features of the affected genes. First, differentially expressed genes were tested for bias towards higher or lower expression, and we found no difference in the distribution of expression values (Figure 3C). Expression breadth across tissues was also assessed, as perhaps differentially expressed genes were biased towards tissue-specific expression. Again, no difference in the distribution of expressed tissues was found (Figure 3D). Next, we considered whether differentially expressed genes had fewer lines belonging to the minor structural haplotype, which might suggest a technical bias influenced by sample size. For this test, the minor structural haplotype was defined as the second most frequent structural haplotype included in the statistical test. Once again, no difference was observed between the distributions when considering frequency distribution across all lines (Figure 3E) or just those included in the statistical test (Figure 3F), indicating no bias towards low frequency structural haplotypes. Finally, we evaluated the evolutionary conservation of these genomes. Pan-genome classifications represent the frequency of a B73 gene across the lines as either core (present in all 26 lines), near-core (present in 24-25 lines), or dispensable (present in 2 – 24 lines). We hypothesized that differentially expressed genes might be overrepresented amongst dispensable genes, but we found no difference in the proportion of genes belonging to any of these classifications (Figure 3G). Zooming out in evolutionary time to the phylostrata distribution of genes, once again, no enrichment of either younger or older phylostrata classifications was observed amongst differentially expressed genes (Figure 3H).

### Differentially Expressed Genes Lack Strong Functional Signatures

To test for functional similarities among the differentially expressed genes, we first performed a Gene Ontology (GO) enrichment analysis. Prior to enrichment testing, we compared the proportion of genes with assigned GO terms between differentially expressed and non-differentially expressed genes as differences could indicate unequal functional characterization and found no significant difference (p = 0.18). We compared the proportion of enriched terms relative to the Promoter-SV gene set and, for both differentially and non-differentially expressed genes, we observed fairly broad enriched terms. The only exception was the enrichment of biological processes related to RNA processing amongst differentially expressed genes, but, generally speaking, we did not observe any specialized biological functions amongst the differentially expressed genes (Figure S6).

To assess potential functional importance more directly, we evaluated the presence of differentially expressed genes among several compiled lists of maize genes that have evidence of being functional genes based on previous studies. This included curated genes, GWAS hits, and candidate domestication genes, all of which represent genes that have either been associated with or experimentally validated for functional importance. Using chi-square tests, we detected no enrichment of domestication genes among differentially expressed genes, but we did observe a significant de-enrichment for curated and GWAS-associated genes (Table 2). These results suggest that differentially expressed genes may have reduced functional importance.

**Table 2.** Chi–Square Tests for Enrichment of Functional Genes Amongst Differentially Expressed Genes.

| Gene List | All Promoter SV<br>vs<br>Differentially<br>Expressed | All Promoter SV<br>vs<br>Non-Differentially<br>Expressed | Differentially Expressed<br>vs<br>Non-Differentially<br>Expressed |
| --- | --- | --- | --- |
| Curated Genes<br>(N = 3,813)<br>% in list | 5.02 x 10<sup>-5</sup> ***<br>12.7% vs 10.2% | 2.31 x 10<sup>-2</sup> *<br>12.7% vs 13.8% | 1.90 x 10<sup>-8</sup> ***<br>10.2% vs 13.8% |
| GWAS Hits<br>(N = 6,666)<br>% in list | 3.16 x 10<sup>-3</sup> **<br>22.7% vs 20.3% | 9.15 x 10<sup>-2</sup><br>22.7% vs 23.7% | 5.87 x 10<sup>-5</sup> ***<br>20.3% vs 23.7% |
| Candidate<br>Domestication Genes<br>(N = 1,213)<br>% in list | 8.66 x 10<sup>-1</sup><br>2.5% vs 2.5% | 9.30 x 10<sup>-1</sup><br>2.5% vs 2.4% | 8.02 x 10<sup>-1</sup><br>2.5% vs 2.4% |

Pearson's Chi-squared test with Yates's continuity correction. \*, \*\*, and \*\*\* refer to a p-value less than 0.05, 0.01, and 0.001 respectively

### TE Content of Promotor SVs is The Only Feature Associated with Different Expression

Given the role of transposable elements (TEs) in generating SVs and their potential to contain regulatory elements and drive chromatin modifications, both of which can impact expression of nearby genes, we evaluated the TE content of promoter SVs. No difference was observed in the raw amount of SV sequence derived by TEs between differentially expressed and non-differentially expressed genes (Figure 4A,B). Similarly, if we looked at the category of the SV defined based on the degree of TE overlap (following Munasinghe et al. 2023a’s classifications), no difference was found between differentially and non-differentially expressed genes for the proportion of promoter SVs described as either ‘No TE SV’, ‘Incomplete TE SV’, ‘TE = SV’, ‘Multi TE SV’, or ‘TE Within SV’. This suggests that the simple presence of TE sequences within promoter SVs is not a reliable predictor of expression differences.

**Figure 4.**
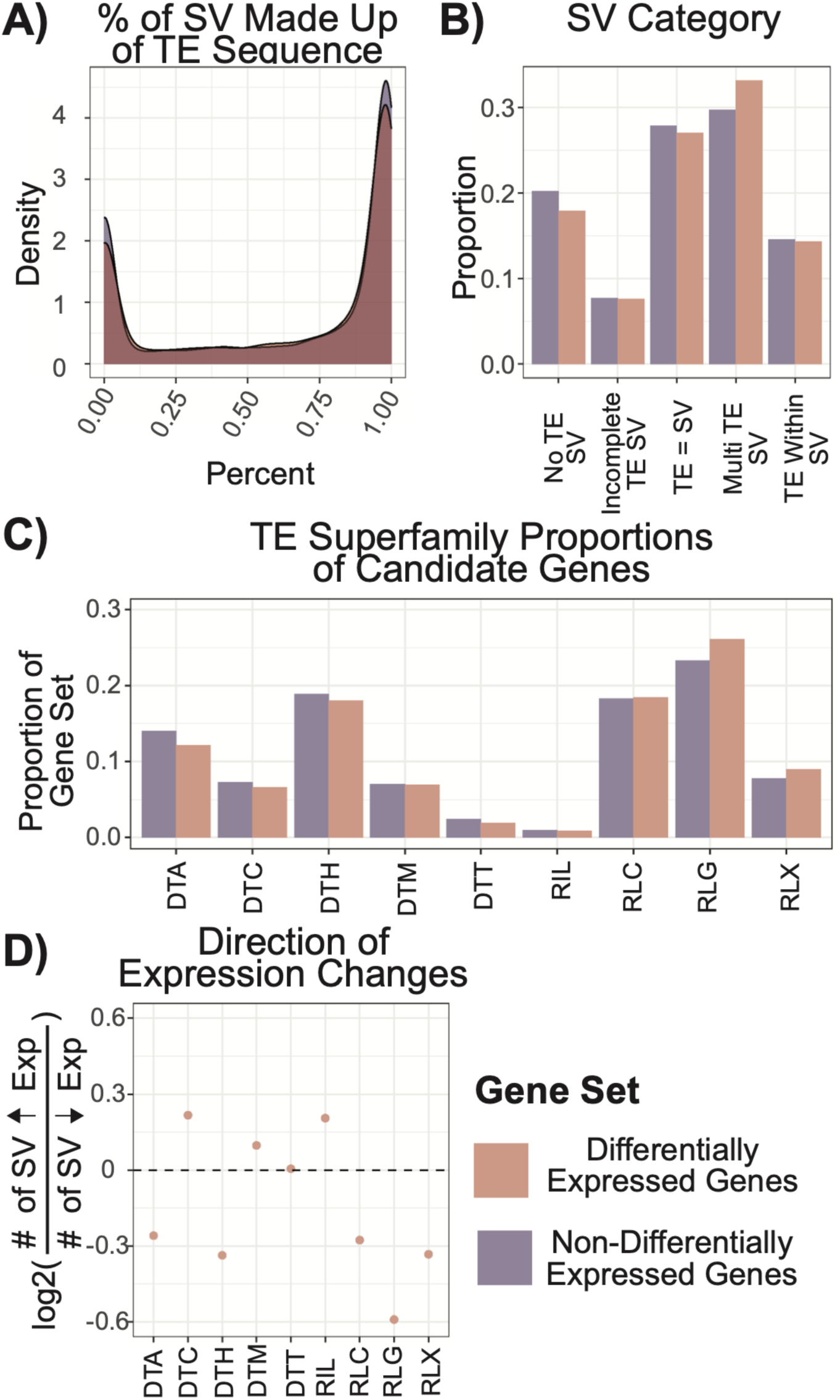
Transposable Element Content is Associated with Differences in Expression Between Structural Haplotypes. We contrasted the content of promoter SVs overlapping TE sequences for differentially (pink) and non-differentially expressed (purple) genes. (A) The distribution of the percent of promoter SVs overlapping TE sequences was contrasted. (B) Munasinghe et al. 2024 categorized each SV based on how it overlapped TE sequence and the proportion of all SVs by category was plotted. (C) The proportion of promoter SVs overlapping each TE superfamily was plotted by whether or not that promoter SV was associated with differential expression. (D) For differentially expressed genes, we took the log2 ratio of the number of genes where presence of the SV sequence resulted in increased versus decreased expression and plotted this by TE superfamily.

A difference emerged when we examined the class and superfamily of TE-mediated promoter SVs. Specifically, retrotransposons, particularly RLGs and RLXs, were enriched amongst promoter SVs associated with differential expression, while DNA transposons, notably DTAs and DTCs, appeared to be de-enriched (Figure 4C). This is intriguing given that DNA transposons, which tend to insert near genes, were initially enriched amongst the Promoter-SV genes. Zooming in to a family-level analysis revealed no statistically significant families.

The presence of a TE can introduce regulatory elements which could potentially increase expression of nearby genes, but the presence of a TE can also result in epigenic silencing, which could potentially decrease expression of nearby genes. To explored whether the presence of a TE-mediated promoter SV resulted in increased or decreased expression of the nearby gene, we focused on the subset of differentially expressed genes with only two structural haplotypes that differed by a single promoter SV (N = 1,304). Insertions containing DNA transposons often had a mixed result, but, generally speaking, they were more often associated with increases in downstream expression. Insertions containing LTR retrotransposons, on the other hand, were consistently associated with reductions in downstream expression (Figure 4D).

Differences in the epigenetic profiles surrounding TE-containing promoter SVs could be contributing to the observed enrichments. In maize, it has been shown that, while H3K9me2 is enriched over both DNA transposons and LTR retrotransposons, LTRs show a greater enrichment of this repressive chromatin mark [50]. These chromatin modifications could impact the accessibility of the gene promoter and potentially explain the observed reductions in expression associated with retrotransposon-containing promoter SVs. To explore this, we examined whether the presence of an SV altered the accessibility of the promoter using previously identified unmethylated regions (UMRs) and accessible chromatin regions (ACRs).

Promoter-SV genes with two structural haplotypes differing by only a single TE-overlapping SV were analyzed, agnostic to whether or not they were differentially expressed (N=4,633). For each gene, we could determine the proportion of lines for each structural haplotype that intersect an accessible feature at the site of the promoter SV and then calculate the difference in this proportion between haplotypes. For UMRs, 77% of genes had no difference in the proportion overlapping between structural haplotypes, while only 50% of genes had no difference in the proportion of overlapping ACRs. Very few genes exhibited a complete difference between structural haplotypes, with 22 and 13 for UMRs and ACRs, respectively. Only 536 genes showed a difference in both overlapping UMRs and ACRs. Regions of accessible chromatin tend to be more variable between tissue types than DNA methylation, which tends to be more stable in plants [51–53]. Consequently, accessible chromatin regions may shift more often than DNA methylation patterns, which may explain the difference in the proportion of overlapping UMRs versus ACRs.

The UMRs and ACRs that were used for this analysis were identified from a single tissue, seedling leaf, and so their differential expression across all tissues may not reflect the relationship between the epigenetic state of the promoter SV and the differential gene expression. As such, we further narrowed in to look at if each gene was differentially expressed specifically in leaf tissue. When focusing on just leaf tissue, we did not observe any differences in the distribution of differences for the proportion of overlapping UMRs and ACRs between differentially expressed and non-differentially expressed genes (asymptotic two-sample KS test p-value 0.14 and 0.11 respectively). While nonsignificant, there is a subtle elevation of differences for both overlapping UMRs and ACRs of differentially expressed genes (Figure S8), but, in totality, our results do not suggest that chromatin accessibility alone explains why TE-mediated promoter SVs are associated with changes in gene expression.

## Discussion

In this study, promoter structural variation was analyzed across 26 diverse maize lines to assess the genome-wide impact of promoter SVs on gene expression. We identified 11,737 genes where, for at least a subset of lines, the only structural differences occurred within the 1kb promoter region upstream of the transcription start site for the canonical transcript. 30% (N=3,584) of these genes exhibited significant differential expression between structural haplotypes, more than fivefold higher than expected by chance. This enrichment was not explained by promoter SNP content, indicating that structural variation is independently associated with expression divergence. Differentially expressed genes more often exhibited tissue-specificity, with 2 – 5 tissues driving the observed expression differences. In addition to mean differences in expression, promoter SVs were also associated with larger differences in expression variance across tissues.

The only distinguishing characteristic of promoter SVs associated with differential expression emerged when their overlap with TE content was explored. The simple presence or absence of TE sequence within a promoter SV was not associated with differential expression. However, differences were observed when the analysis was done by TE class. LTR retrotransposons were enriched amongst promoter SVs associated with differential expression and generally resulted in reduced expression. DNA transposons, on the other hand, were less frequently associated with differential expression, but, when they were, they were generally correlated with increased expression of the downstream gene. Functional differences between class I and class II transposons may explain these differences. Class I retrotransposons are highly prolific, as they leverage a ‘copy-and-paste’ mechanism for transposition where they are first transcribed into an RNA intermediate which is the reverse-transcribed and re-integrated into new genomic positions [54–56]. DNA transposons have lower transposition rates as they leverage a ‘copy-and-paste’ mechanism where the TE sequence is excised and then reintegrated elsewhere within the genome. While both classes contain key sequences required for their transposition, they still rely on the host’s cellular machinery to generate relevant mRNA transcripts. TEs consequently harbor transcription factor binding sites, enhancers, and promoters in order to exploit host transcription and increase their insertion rate. An unintended consequence is that these features may also increase the expression of nearby genes. Host silencing mechanisms have evolved to target these internal regulatory sequences in order to prevent TE activity, though their deposition and maintenance is imperfect and can spread into nearby sequences resulting in reduced gene expression.

Due to their highly proliferative nature, LTR retrotransposons are often hyper-methylated relative to DNA transposons and frequently co-localize with repressive histone modifications [50,57,58]. This might explain why LTR retrotransposons are more often associated with changes, notably reductions, in expression. Interestingly, DNA transposons, while less proliferative, tend to preferentially insert into accessible regions upstream of genes [59–61]. By inserting into accessible regions, the regulatory features housed within DNA transposons may be exposed which explains why they were sometimes associated with increases in expression. It is important to note that these trends represent enrichments and that both classes exhibit associations with genes showing both increased or decreased expression. While there may be broad patterns delineating LTR retrotransposons from DNA transposons, it is clear that TE-mediated promoter SVs can change genome-wide expression patterns in complex and unpredictable ways.

Our results align with examples derived from single-locus characterizations. Within maize, a prime example of this phenomenon is the expression of the domestication gene *tb1*. *tb1* is a major contributor to the increased apical dominance during maize domestication, and research has shown that the maize allele of *tb1* is much more highly expressed relative to the ancestral allele in Mexican teosinte [62]. This expression difference is due to the insertion of a *Hopscotch* retrotransposon upstream of *tb1*. In cultivated grape (*Vitis vinifera L.*), the insertion of a retroelement into the promoter of the gene *VvMybA1* is responsible for white wine grape varietals [63–65]. Epigenetic silencing via DNA methylation of this retroelement is associated with reduced expression of *VvMybA1* causing the white berry phenotype. Demethylation of this region allows for higher expression which causes color reversions. In Arabidopsis, a SINE DNA transposon immediately upstream of the *FLOWERING WAGENINGEN* (*FWA*) locus causes the epigenetically silencing of this gene in vegetative tissues [66]. In mutants, a stable epimutation can emerge where ectopic expression of *FWA* leads to a late flowering phenotype. From these, and other, examples, it is clear that the epigenetic silencing of TEs can change downstream gene expression patterns, but our results suggest these functional examples may be infrequent across the genome.

Combined, this evidence supports an interesting evolutionary and regulatory paradigm. The epigenetic silencing of TEs may cause the downregulation of adjacent genes. However, we know that epigenetic modifications shift in response to environmental stimuli. This rewiring of the epigenome could mask or expose the internal regulatory sequences within TEs allowing for context-dependent changes in gene expression [67]. However, it is unclear whether this is truly occurring broadly across the genome. Most changes in gene expression are expected to be deleterious, and, if TEs influence expression in this manner, they will likely experience purifying selection. On the other hand, if this change is truly adaptive, we expect it to fix within the population. Consequently, segregating TE variation (which is the only type we can test in our analysis) likely persists in the genome because it is (1) selectively neutral and has no phenotypic effect or is (2) under balancing selection due to environmental fluctuations or other selective pressures. The fact that differentially expressed genes show very little functional signatures and exhibit substantial tissue-, or context-, specificity support both of these explanations.

While our analysis reveals novel evidence that TE-mediated promoter SVs induce expression changes across the genome, there are several technical limitations that impede our ability to fully investigate this phenomenon. Our statistical analysis required variants to be segregating at moderate frequency, which, given the effects of selection, likely limit our ability to identify variants which have large phenotypic effects. Also, while we hypothesized that differences in epigenetic silencing may explain the different outcomes observed between LTR retrotransposons and DNA transposons, we did not find any differential overlap with unmethylated regions or accessible chromatin regions. Our results were very sensitive to the tissue, or context, tested, and it is very likely there are tissue-specific differences in the epigenome we were unable to capture. Therefore, it remains unclear the extent to which changes in the epigenome are able to explain the results of this study. This is likely also influencing the genes we are able to identify as differentially expressed. Exposure of these lines to alternative environmental contexts (e.g. abiotic or biotic stress) would likely reveal context-dependent differential expression among genes we currently classify as non-differentially expressed and may even uncover genes with functional roles not captured by existing annotations. A phenotype-driven experimental design that targets specific traits of interest may be better suited for linking specific TE-mediated variation with adaptive differential expression.

## Conclusions

Our analysis shows widespread, tissue-specific differences in expression associated with TE-linked promoter structural variants within maize. Our analysis focused on segregating variation and identified key differences in how LTR retrotransposons and DNA transposons may change the expression of nearby genes. Most genomes contain substantially less TE content, and there is even observable differences in host silencing of TEs between maize and Arabidopsis (a plant model species with substantially less TE content). Consequently, the potential of TEs to regulate gene expression is likely defined by the unique co-evolutionary relationship between TE content and host silencing within a species. However, the findings and patterns identified within maize may be translatable into other plant systems. Plastic gene regulation is crucial for plant survival, as they cannot simply move to a new location in response to environmental stress. TEs provide a medium through which host genomes can induce context-specific gene expression, and it will be vital to further explore this relationship to engineer resilient crop species.

## Methods

### Genome Alignments and Transposable Element Annotations

Pairwise whole-genome alignments between the B73 reference genome assembly and each of the other NAM inbred founder lines assemblies [47] were generated using AnchorWave [68] as described in Munasinghe et al. [49]. In that study, TE annotations for each genome that were generated using panEDTA [69] were filtered to remove non-TE repeats, helitrons, and elements exhibiting features indicative of misannotation. For the present analysis, we utilized both the summarized pairwise genome alignments and the filtered TE annotations, which are publicly available on Dryad (https://datadryad.org/dataset/doi:10.5061/dryad.5qfttdz9t).

### B73 Gene Feature Information

To obtain B73 gene information, we downloaded the previously generated genomic feature file (gff3) containing the B73 gene annotation (https://www.maizegdb.org/genome/assembly/Zm-B73-REFERENCE-NAM-5.0). Genes located on a scaffold were filtered out, and only the canonical transcript was retained for the remaining genes. This left us with a total of 39,035 B73 gene annotations. Each gene was divided into three distinct features - the promoter, exonic sequences, and intronic sequences. Exonic sequences included annotated UTRs when available. When defining the promoter, we considered two different thresholds - both 1kb and 2kb upstream of the transcription start site (defined as the start of the first exon). Results were very similar between the 1kb and 2kb analysis, and, as such, the 2kb results are only presented in the supplemental materials.

### Transcriptomic Data Across 10 Maize Tissues

Raw RNA-sequencing data released with Huffored et al. 2021 (https://www.ebi.ac.uk/ena/browser/view/ERX3793507) [47] was downloaded for each line. RNA-sequencing was performed across ten different tissues: root, shoot, leaf base, leaf, leaf tip, tassel, ear, anther, endosperm, and embryo. The number of biological replicates per tissue ranges from zero to four across these datasets with most having three biological replicates. The number of replicates for some tissues, notably endosperm and embryo, were zero due to difficulties extracting the necessary tissue.

### Epigenomic Landscape for Each Line

Unmethylated regions (UMRs) and accessible chromatin regions (ACRs) were identified for each of the lines using Enzymatic Methyl-sequencing and ATAC-sequencing peaks respectively generated from seedling leaf tissue [47]. Locations of UMRs (https://ars-usda.app.box.com/v/maizegdb-public/folder/176807408173) and ATAC ACRs (https://ars-usda.app.box.com/v/maizegdb-public/folder/165367078135) for all lines were downloaded.

### Curated Maize Gene Lists

Previously constructed lists of maize genes associated with different functions were downloaded. Brohammer et al. 2018 collated curated genes (derived from Schnable and Freeling 2011) and GWAS hits (derived from Wallace et al. 2014). Both of these gene lists represent genes that have either been associated with or functionally validated for functional impact. These gene lists were downloaded at https://github.com/HirschLabUMN/maize-fractionation/tree/master/data/assests. Hufford et al. (2012) constructed a list of candidate genes associated with evidence for selection during domestication. This gene list was provided with the supplementary materials for this analysis (Supplementary Table 6 in Hufford et al. 2012) [70]. All of these gene lists were originally made for prior versions of the reference genome (B73 v4 for curated genes and GWAS hits, B73 v3 for domestication genes) and coordinate lift overs were identified using MaizeGDB (https://www.maizegdb.org/gene_center/gene - “Translate Gene Model IDs”). This led to 3,813 curated genes, 6,666 GWAS hits, and 1,213 domestication genes.

### Generating Structural Haplotypes for B73 genes

Structural variants were identified from summarized pairwise whole-genome alignments between the B73 reference genome and every other line. An SV was defined as a contiguous sequence of > 50 base pairs that was present in one genotype but completely absent in the other. By definition, each SV had a distinct start and end position in the genotype where the sequence was present and a single start and end position in the genotype where it was absent. For each SV, we extracted its unique identifier, chromosome, and coordinate positions relative to the B73 reference genome assembly.

To determine whether an SV in one pairwise comparison corresponded to an SV in another, we defined two SVs as the same SV if all of the following criteria were met: 1) the start positions were within 50 base pairs of each other, 2) the end positions were within 50 base pairs of each other, 3) the size of the SV differed by less than 100 base pairs, and 4) the category (assigned according to Munasinghe et al. 2023a) was the same (Figure S1). Categories from Munasinghe et al. (2023a) were based on overlaps with TE annotations and included: ‘No TE SV’, ‘Incomplete TE SV’, ‘TE = SV’, ‘Multi TE SV’, and ‘TE Within SV’. SVs that met these criteria were assigned a unique numerical ID, which allowed them to be linked across pairwise alignments.

We focused on annotated chromosomal B73 genes and, for each gene, defined three distinct features based on the canonical transcript of that gene - the gene promoter, exonic sequence, and intronic sequence. We defined the promoter as either 1kb or 2kb upstream of the transcription start site. Using bedtools (v.2.29.2), we intersected the coordinates (in B73) of every SV with these gene features. Comparable SVs overlapping a specific feature were assigned a two-character string identifier (Figure S1): the first character indicated the overlapping feature (P = promoter, E = exon, I = intron) and the second character was a numerical tag distinguishing different SVs overlapping the same feature.

By concatenating these identifiers across all gene features, the structural haplotypes were generated for each gene across all the lines. B73, as the reference in all pairwise comparisons, was assigned the numeric tag 0. Any line without SVs relative to B73 shared this feature. For example, a gene in a given line with no SVs in any feature would have the structural haplotype P0_E0_I0, whereas a haplotype P1_E0_I0 indicates a single SV in the promoter but no SVs in exons or introns. Importantly, these structural haplotypes capture only structural variants; lines sharing the same structural haplotype may still differ at the SNP or small InDel level.

### Processing of Raw Gene Expression Data

Although gene annotations exist for all 26 lines, the B73 v5 reference gene annotations represents the highest-quality annotations in the population. Consequently, gene features for B73 genes were the most robust and accurate, making it the preferred reference for downstream analyses. As such, raw RNA-sequencing reads from each line were mapped to the B73 v5 reference genome. The reference genome was first indexed using STAR (v2.5.3a)[71]. Reads were then trimmed using Trim_Galore (v0.4.3) [72] and subsequently aligned to the B73 reference genome using STAR. Gene-level counts were generated with HTSeq (v0.7.2) [73] and normalized in R using DESeq2 (v.1.48.1) [74].

### Identification of Promoter-SV Defined Genes for Differential Expression Analyses

To test whether structural variation in the promoter region upstream of a gene is associated with differential expression, we first identified genes with at least two distinct structural haplotypes, each represented by a minimum of three lines. We only considered structural haplotypes with no structural variation within the gene body (i.e., haplotypes classified as E0_I0) to ensure that structural differences were confined to the promoter region. Additionally, as RNA sequencing reads were mapped to the B73 reference genome, this minimizes technical artefacts due to mismapping caused by genic structural variants.

Differential expression analyses for a given gene included only the lines belonging to haplotypes meeting these criteria. Given this, the minimum number of lines per tested gene was six (two structural haplotypes with three members each). To remove non-expressed genes, we employed a minimum expression threshold of at least one normalized count greater than four across all expression values. This filtering yielded 11,737 genes representing approximately 30% of the original gene set. We refer to this set of genes as the Promoter-SV gene set as they are characterized by segregating structural variation within the promoter and meet the minimum requirements necessary to test for differential expression.

### Testing for Differential Expression Between Structural Haplotypes

Promoter-SV genes differ in which lines were included, depending on the structural haplotypes present for each gene. Consequently, differential expression analysis was performed on a gene-by-gene basis. We used the following mixed-effects linear model:

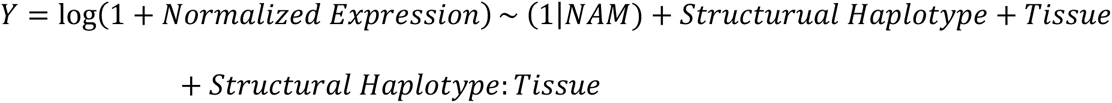

In this model, structural haplotype, tissue, and their interaction were treated as fixed effects, while the line was treated as a random effect to account for replicate measurements. Raw p-values for each term were obtained using a Type II analysis of variance.

Although the same model was applied across all Promoter-SV genes, the composition of each group varies by gene. This prevents the use of traditional multiple testing corrections. Instead, a permutation-based approach was implemented. For each gene, expression values were shuffled among lines within each tissue (i.e., values remained within tissues but were randomized across lines and consequently across structural haplotypes). Five permuted datasets were generated and analyzed using the same mixed-effects model (Figure S8). A gene was considered significantly differentially expressed between structural haplotypes if its p-value for the structural haplotype term in the true data set was < 0.05 and it was not significant (p >= 0.05) in at least four of five permuted data sets.

### Generation of Random and SNP-Based Haplotypes

To contextualize the results of our analysis, we generated both a Promoter-Random gene set and a Promoter-SNP gene set. Our Promoter-SV gene set contains 11,737 candidate genes, each associated with two or more structural haplotypes with a subset of lines assigned to each haplotype. To generate the Promoter-Random gene set, we randomly sampled 11,737 non-tested B73 genes, and, for each gene, we copied the haplotype configuration of a corresponding Promoter-SV gene. For each randomly sampled gene, only the lines present in that corresponding Promoter-SV gene were included and assigned to the matching structural haplotypes. This allowed us to copy the haplotype structure and line composition for each Promoter-SV gene and apply it to a random counterpart, while keeping the original expression values of the random counterpart. If a random gene did not meet our expression requirements, it was dropped and another gene was randomly chosen. The same differential expression tests were run as described above using this Promoter-Random gene set, and we compared results from our true Promoter-SV gene set to those found under the constructed Promoter-Random gene set.

We also investigated whether our results were influenced by single nucleotide polymorphism (SNP) content within the promoter regions. To do this, we constructed SNP haplotypes for the 1kb promoter by extracting all SNPs overlapping that region for each gene from the pairwise whole-genome alignments. These SNPs were concatenated and used in place of the promoter structural haplotype, while the exonic and intronic structural haplotypes were retained. Thus, the only difference between the structural and SNP haplotypes for a given gene was the definition of the promoter haplotype - based on SV or SNP content, respectively. Candidate genes for the SNP haplotype analysis were defined using the same criteria as for the structural haplotypes: no SVs overlapping the gene body, at least two SNP haplotypes with a minimum of three lines each, and at least one expression value exceeding four normalized counts. This yielded 10,576 genes referred to as the Promoter-SNP gene set. The same statistical model as described above was used to test for differential expression, replacing the structural haplotype term with a SNP haplotype term.

### Tissue-Specific Tests of Differential Expression

For the previous ten-tissue analysis, we did not permute expression values across tissue when identifying significant differentially expressed genes. We, therefore, could not use that model to assess tissue-specificity. Instead, we repeated our analysis tissue by tissue and adjusted our model by simply dropping terms containing tissue as a fixed effect. This allowed genes to be identified that were differentially expressed within each tissue. A Promoter-SV gene may not be expressed in a given tissue, as our expression requirements for inclusion simply required one value be greater than four normalized counts. In instances when a tissue had no expression across all included genotypes, it was not tested. The vast majority of genes (N =9,111 out of 11,737, ∼78%) were tested in all ten tissue-specific analyses.

### Identifying Outliers for Expression Variability Across Tissues

To test whether expression variance across tissues differed between structural haplotypes the coefficient of variation and the mean absolute deviation were tested on the subset of structural haplotypes that differed by only a single SV (N=5,872). The coefficient of variation was calculated as the standard deviation in expression across tissues divided by the mean and represents the relative variability in expression. The mean absolute deviation was calculated as the average distance between each expression value and the mean and represents the relative spread of the data. These two metrics are analogous but explored slightly different features of expression variance across tissues. For both variance metrics, the mean value was calculated for each structural haplotype belonging to a given gene and the absolute difference between the values was computed for the two structural haplotypes. Genes in the bottom 2.5% of the distribution of values represented those where the expression variance was nearly identical between both structural haplotypes, while genes in the top 2.5% represented those where the two structural haplotypes exhibited highly divergent expression variance.

### Gene Ontology Analysis

Gene ontology analyses were performed using topGO (v.2.50.0) [75]. PANZZER GO annotations for the B73 reference genome were downloaded from MaizeGDB (https://download.maizegdb.org/GeneFunction_and_Expression/Pannzer_GO_Terms/). We filtered the full list of maize genes to exclude low confidence annotations by removing gene_IDs where the ARGOT score assigned by PANZZER was less than 0.5. We used the “elim” algorithm, as it accounts for GO topology, to perform an enrichment analysis that tests whether there are any overrepresented GO terms within the tested gene set. The enrichment score was calculated using a Fisher’s exact test. Both differentially and non-differentially expressed genes were tested for enriched GO terms. Additionally, genes with high or low expression variability were also tested for enriched terms.

### Enrichment Test of Previously Identified Functional Genes

Previous publications have collated three distinct gene sets that represented various dimensions of functionality. Brohammer et al. 2018 developed both a curated gene list, which included both classical maize genes and those manually curated by the Maize Genetics and Genomics Database (MaizeGDB), and a list of maize GWAS hits, which included genes with GWAS hits across 41 agronomic traits. Hufford et al. 2012 similarly collated a list of domestication genes associated with evidence for selection during domestication. These three gene lists represent maize genes that have either been associated with or experimentally validated for functional importance. As all of these gene lists were originally made for older version of the B73 reference genome, gene_IDs were converted to the B73 v5 gene_IDs using MaizeGDB (https://www.maizegdb.org/gene_center/gene - ‘Translate Gene Model IDs’).

A series of chi-square tests were performed to ascertain whether the presence of functional genes varied for differentially expressed genes. We contrasted differentially expressed genes relative to the full gene set, non-differentially expressed genes relative to the full gene set, and differentially expressed genes relative to non-differentially expressed genes. Differentially expressed genes were considered to have enrichment or de-enrichment, depending on the relative proportions, of functional genes if we observed significant differences for the chi-square tests including differentially expressed genes but not for the test contrasting non-differentially expressed with the full gene set.

### Identifying Whether Presence of SV Sequence Alters Chromatin Accessibility

To explore whether the presence of SV sequence potentially altered the accessibility of the promoter, a subset of Promoter-SV genes was identified that had only two tested structural haplotypes that differ by only a single TE-mediated SV agnostic of whether that gene is differentially expressed (N=4,633). In this set, one structural haplotype was the ‘B73-like’ state, while the other had either an insertion or deletion relative to B73 depending on the type of SV located within the promoter. For each promoter SV, we took at most the 500bp flanking each SV in each genotype, while restricting the flanking regions such that they did not overlap the gene body. Genotypes containing the SV sequence had two distinct flanking regions, one to the left and one to the right of the SV, while genotypes that lacked the SV had a contiguous stretch of at most 1000bp. In rare cases were the flanking regions intersected unalignable or SV sequence the ported coordinates could have the start of the flanking region be larger than the end. In these cases, the lines were dropped from comparison.

To determine whether presence of the SV altered accessibility, the flanking regions were intersected with both the coordinates of unmethylated regions determined via Enzymatic Methyl-sequencing and accessible chromatin regions determined via ATAC-sequencing peaks. For each structural haplotype, the proportion of lines that overlapped these identified features were determined, and the difference between structural haplotypes was determined. It is worth mentioning that most genes in our analysis overlap a feature of accessible DNA, which is unsurprising as these regions were located in the promoter of genes that had some expression.

## Declarations

### Ethics Approval and Consent to Participate

Not Applicable

### Consent for Publication

Not Applicable

### Availability of Data and Materials

The location of all publicly available datasets used in this study has been provided in the methods. All code used to generate the analyses in this study is available on GitHub: https://github.com/HirschLabUMN/NAM_SV_DifferentialExpression. All data sets generated as part of this project are available on Dryad.

### Competing Interests

All authors declare that they have no competing interests.

### Funding

This work was supported by a National Sciences Foundation Grant IOS-1934384 and Postdoctoral Research Fellowship In Biology Award #2410349. This work was also funded by the Minnesota Agricultural Experiment Station.

### Author’s Contributions

M.M., A.R., N.S., and C.H. conceived the project and designed the study. M.M. spearheaded and performed all data analyses. M.M., Y.B., and C.H. designed the statistical framework used to test for differential expression. A.R. re-aligned the RNA-sequencing data to the B73 maize reference genome, while A.S. contributed to the analysis of functional signatures amongst differentially expressed genes. M.M. and C.H. wrote the manuscript, and all authors provided critical revision and input. All authors read and approved the final manuscript.

## Supporting information

Munasinghe_etal_2026_Supplement

## Acknowledgments

We would like to thank Jeff Ross-Ibarra and Beibei Liu for their valuable comments, insights, and suggestions throughout the analysis. We would also like to thank the Minnesota Supercomputing Institute at the University of Minnesota (https://www.msi.umn.edu) for providing resources that contributed to the research results reported within this article.

## Notes

https://github.com/mam737/NAM_SV_DifferentialExpression

### Competing Interest Statement

The authors have declared no competing interest.

