## Supplementary material for "Promoter Structural Variants are Drivers of Genome-Wide Differential Expression in Maize": Munasinghe_etal_2026_Supplement

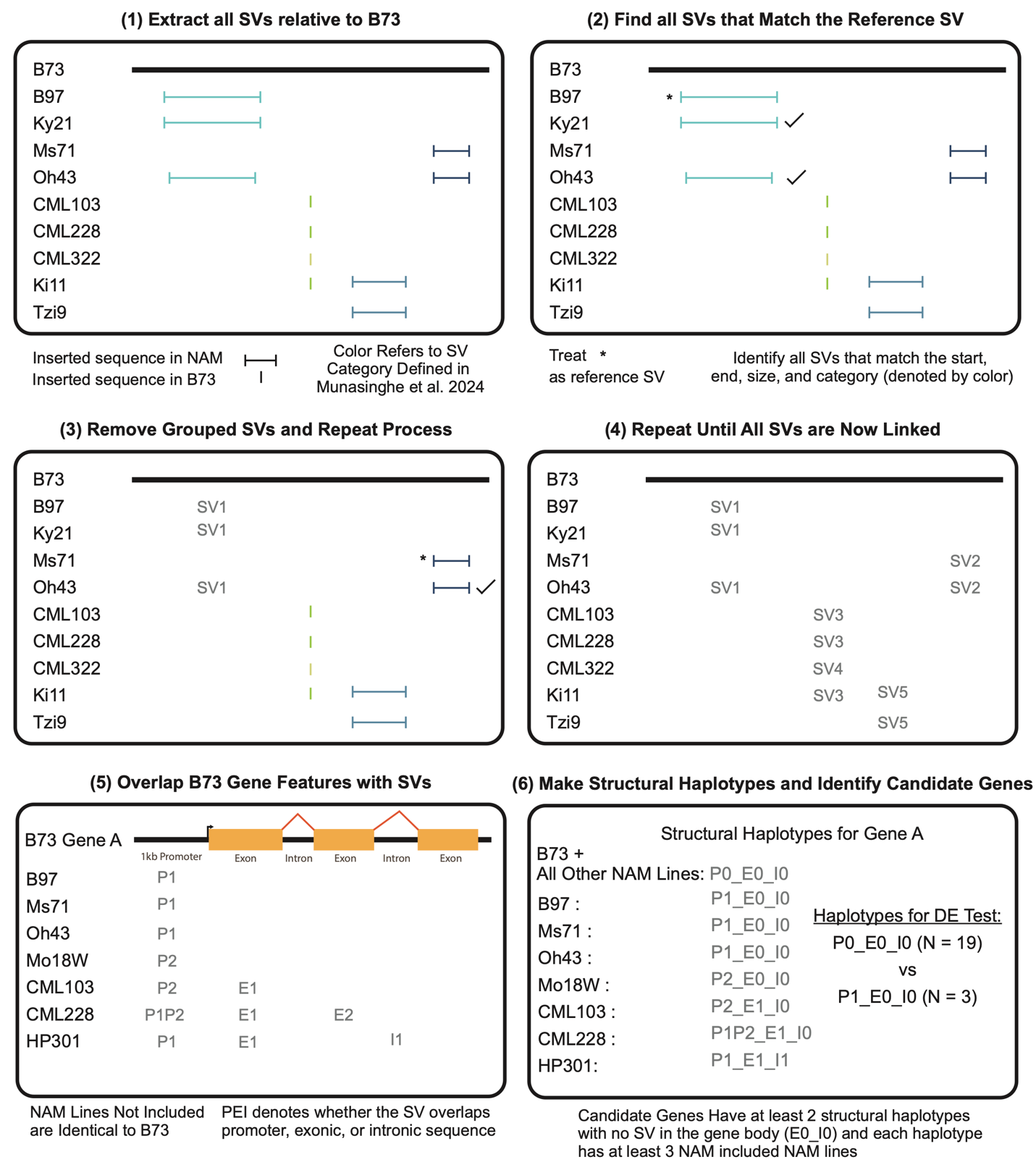

**Figure S1. Visual Schematic of SV Linkage and Structural Haplotype Construction**. Structural variants were identified from pairwise whole-genome alignments between the B73 reference genome and every other NAM founder line. Insertions in B73 relative to a NAM line are visualized by lines with a distinct start and end coordinate, while deletions in B73 relative to a NAM line are visualized by a single vertical bar. We defined SVs as comparable if their coordinate position, size, and overlap with TE sequence (represented by the color of the SV) matched. We could then intersect SVs with B73 gene features to generate structural haplotypes.

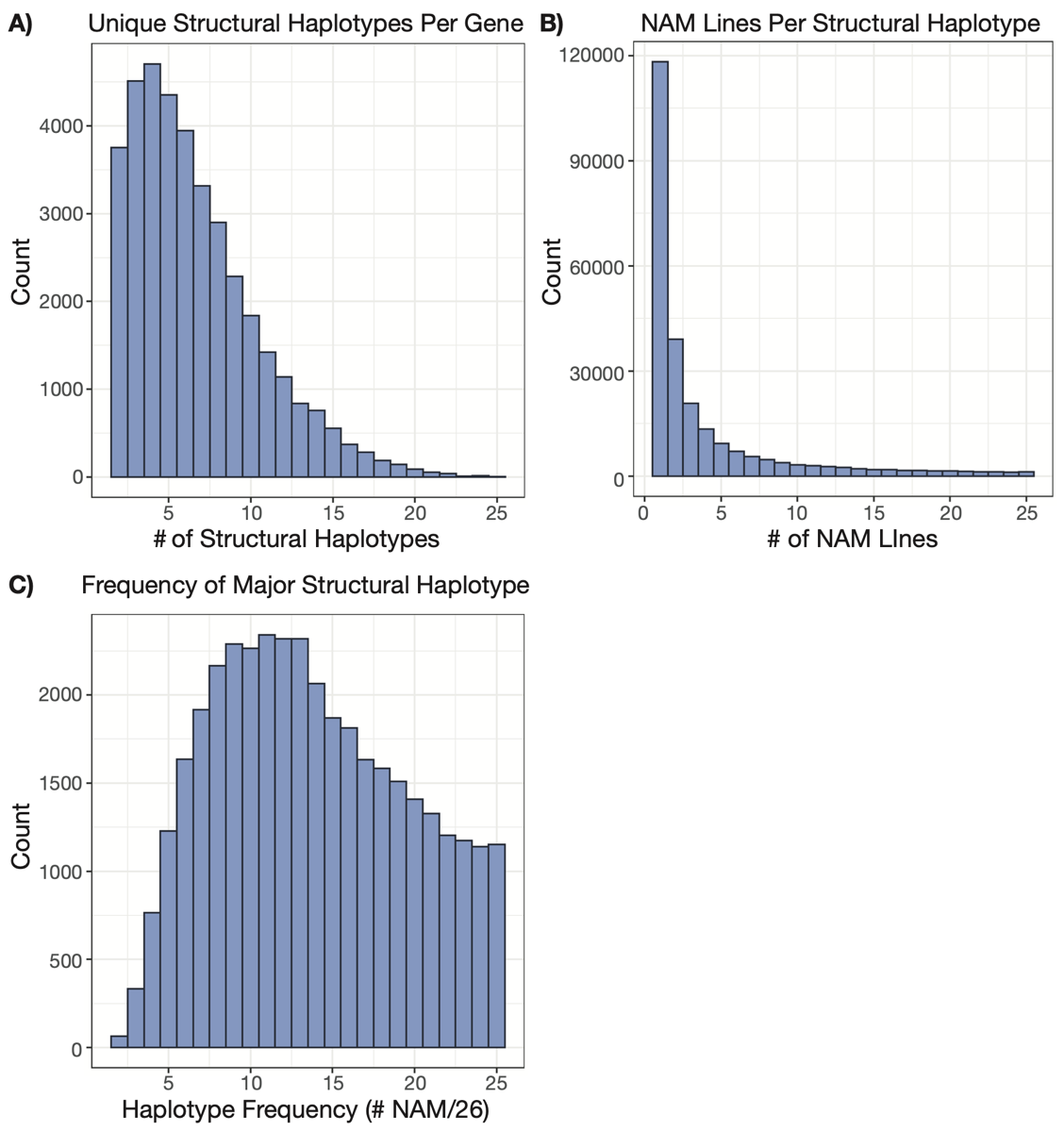

**Figure S2. Structural Haplotypes Across All B73 Genes**. Structural haplotypes for the canonical transcript of all chromosomal B73 genes (N=39,035) were generated. We show distributions of (A) the number of structural haplotypes per gene, (B) how many NAM lines belonged to each structural haplotype, and (C) the frequency of the major structural haplotype of each gene across the NAM founder lines.

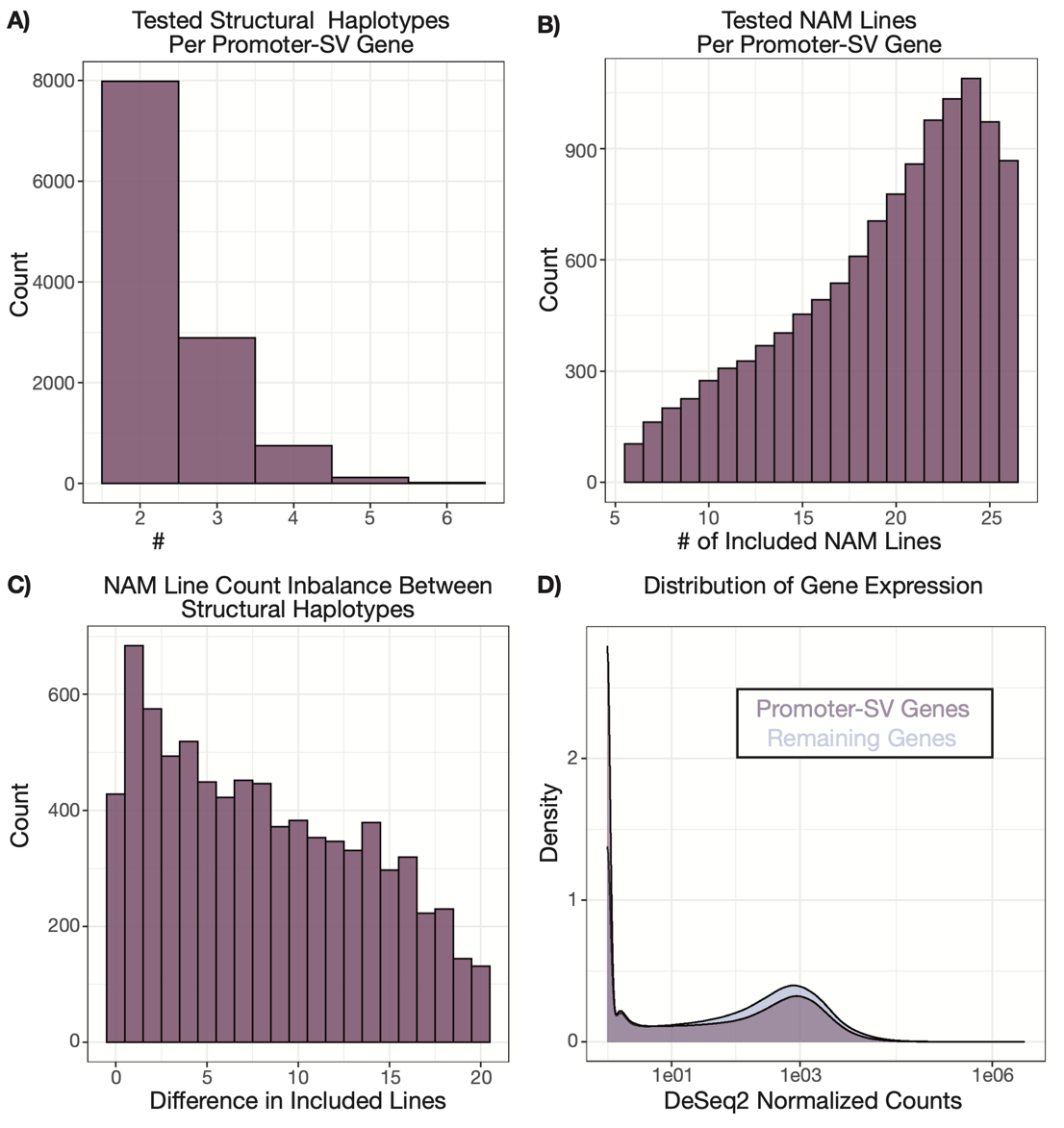

**Figure S3. Structural Haplotypes Across Candidate Genes.** For Promoter-SV genes included in our statistical analysis we show distributions of (A) how many structural haplotypes were tested for each gene, (B) how many NAM lines were included in the test for each gene, (C) how balanced the number of NAM lines is for genes with only two tested structural haplotypes, and (D) normalized expression counts relative to the remaining genes (i.e., those not included in our statistical analysis).

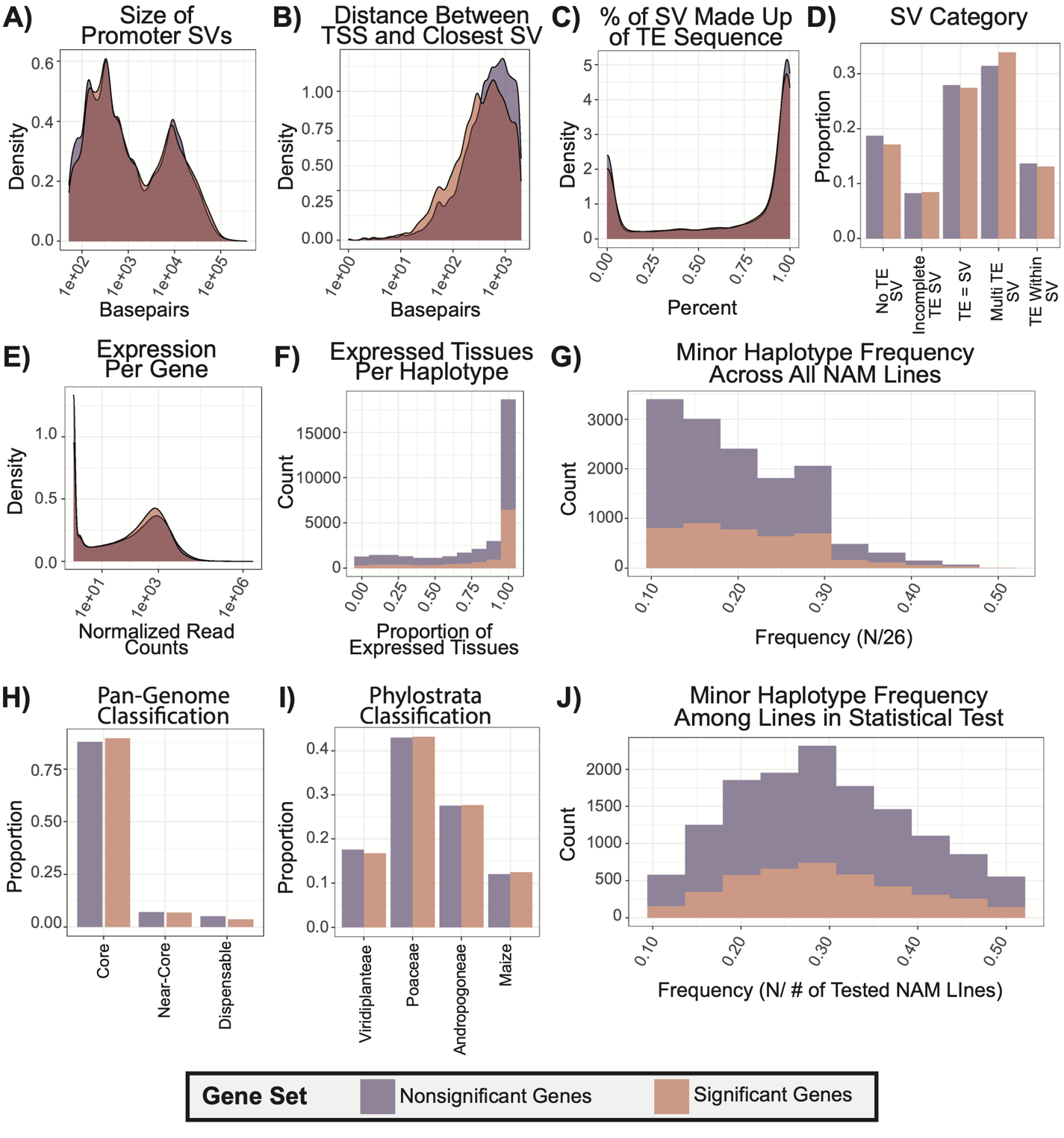

**Figure S4. Limited Differences Between Differentially Expressed and Non-Differentially Expressed 2kb Promoter-SV Genes.** Several different features were contrasted between significant and nonsignificant gene sets including (A) the size of the structural variant located within the promoter, (B) the distance between the transcription start site (TSS) and the closest promoter structural variant, (C) the percent of the promoter structural variant made up of TE sequence, (D) the category as defined in Munasinghe et al. 2024 of the promoter structural variant, (E) the distribution of gene expression values across tissues, (F) the number of expressed tissues per structural haplotype, (G) the allele frequency distribution of the minor haplotype, defined as the second most common haplotype, amongst the entirety of the NAM population, (H) the proportion of each gene set belonging to distinct pan-genome classifications including core (gene is present amongst all 26 NAM lines), near-core (gene is present amongst 23 - 24 NAM lines), or dispensable (shared in 2 - 22 NAM lines), (I) the proportion of each gene set belonging to different phylostrata moving from deeper within the clade (Viridiplanteae) to younger Maize-specific genes, and (J) the allele frequency distribution of the minor haplotype amongst NAM line including in the statistical test of differential expression.

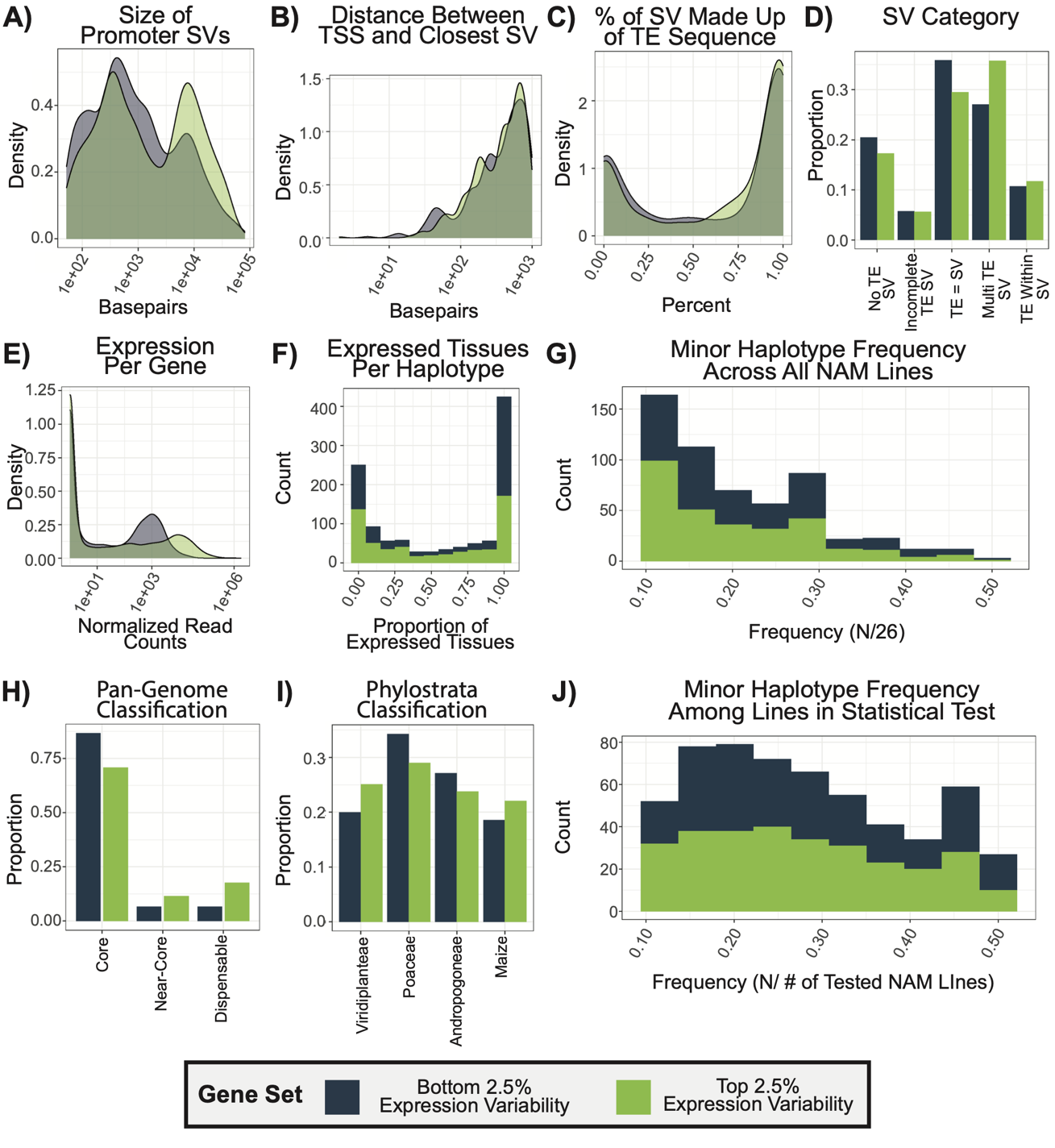

**Figure S5. Differences Between Genes with Low and High Expression Variability Between Structural Haplotypes.** The absolute difference in expression variance was calculated between structural haplotypes, and outliers in the bottom (dark blue) and top 2.5% (light green) were identified. Several different features were contrasted between these genes including (A) the size of the structural variant located within the promoter, (B) the distance between the transcription start site (TSS) and the closest promoter structural variant, (C) the percent of the promoter structural variant made up of TE sequence, (D) the category as defined in Munasinghe et al. 2024 of the promoter structural variant, (E) the distribution of gene expression values across tissues, (F) the number of expressed tissues per structural haplotype, (G) the allele frequency distribution of the minor haplotype, defined as the second most common haplotype, amongst the entirety of the NAM population, (H) the proportion of each gene set belonging to distinct pan-genome classifications including core (gene is present amongst all 26 NAM lines), near-core (gene is present amongst 23 - 24 NAM lines), or dispensable (shared in 2 - 22 NAM lines), (I) the proportion of each gene set belonging to different phylostrata moving from deeper within the clade (Viridiplanteae) to younger Maize-specific genes, and (J) the allele frequency distribution of the minor haplotype amongst NAM line including in the statistical test of differential expression.

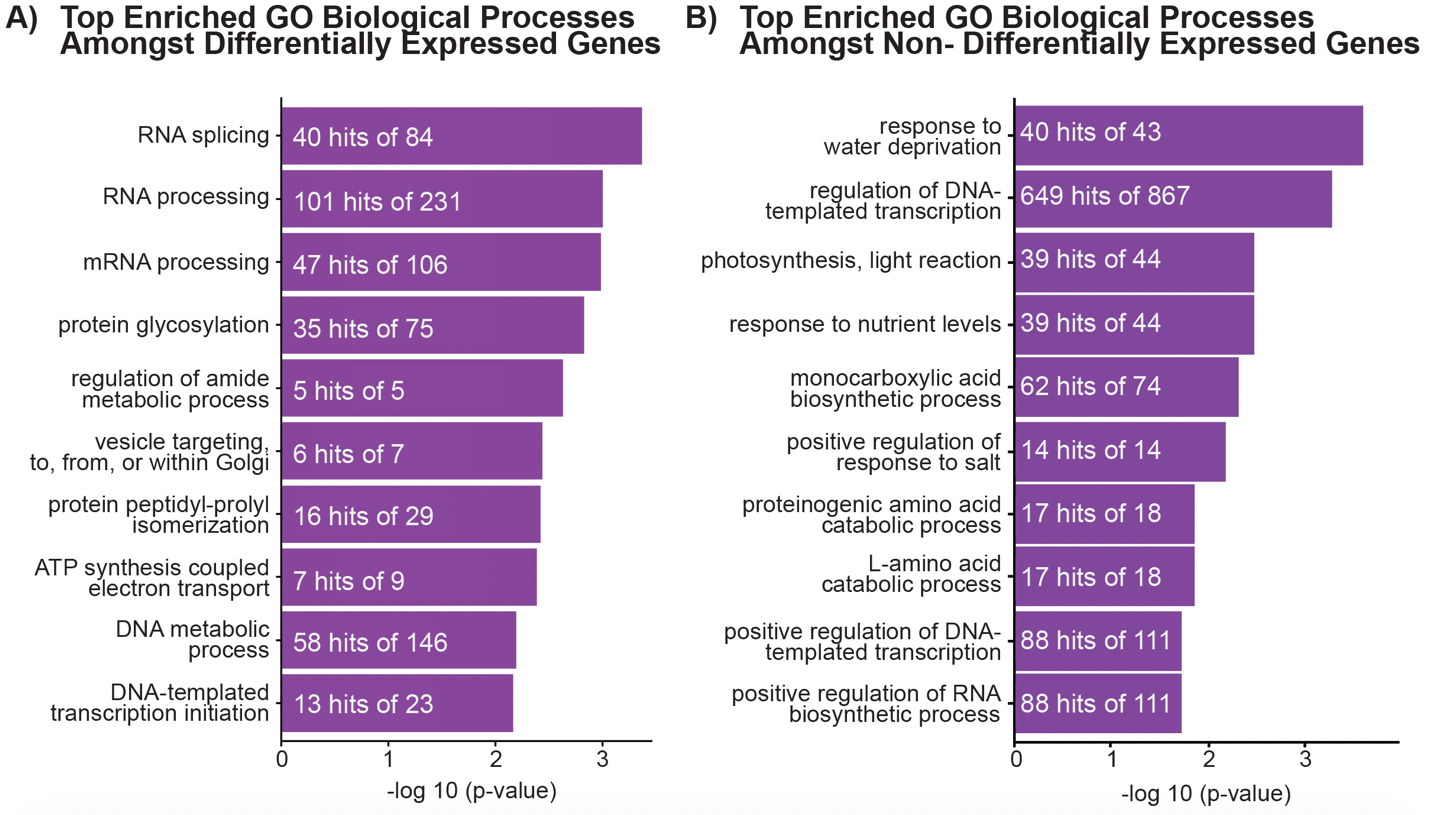

**Figure S6. Enriched GO Biological Processes Amongst Gene Sets**. Enriched GO biological processes were identified for differentially expressed (A) and non-differentially expressed (B) genes relative to the full Promoter-SV Gene Set. The top 10 enriched processes were plotted.

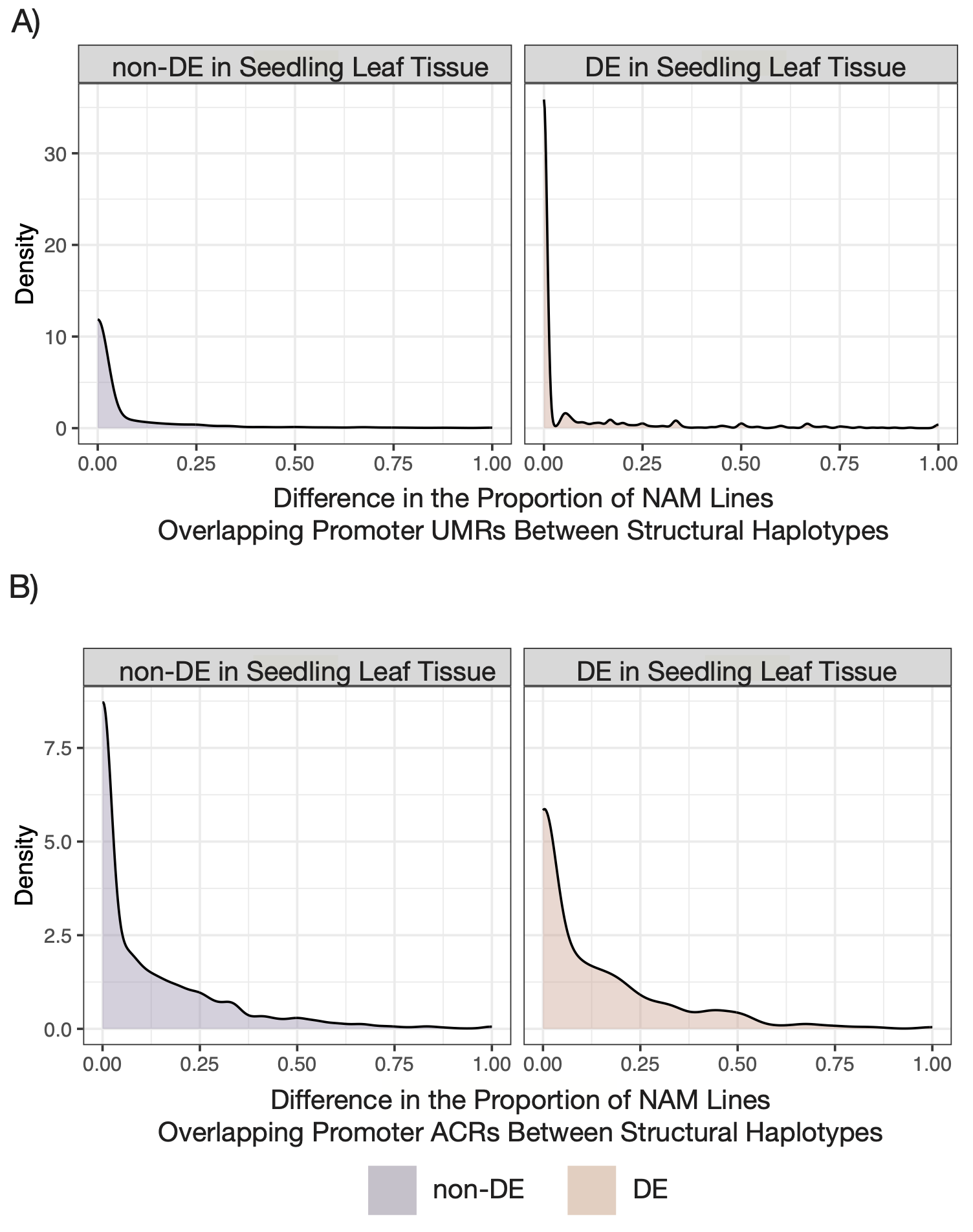

**Figure S7. Distributions of the Difference in the Proportion of NAM Lines Overlapping Regions of Chromatin Accessibility Between Structural Haplotypes**. For Promoter-SV genes with two structural haplotypes that differ by only a single SV (N = 5,872), we determined the proportion of NAM lines for each structural haplotype (presence vs absence of the SV) that intersect an accessible feature and calculate the difference in this proportion between structural haplotypes. (A) Shows the distributions for overlapping unmethylated regions (UMRs) facetted by whether the genes were differentially expressed in seedling leaf tissue. (B) Shows the distributions for overlapping accessible chromatin regions (ACRs) similarly facetted by whether the genes were differentially expressed in seedling leaf tissue.

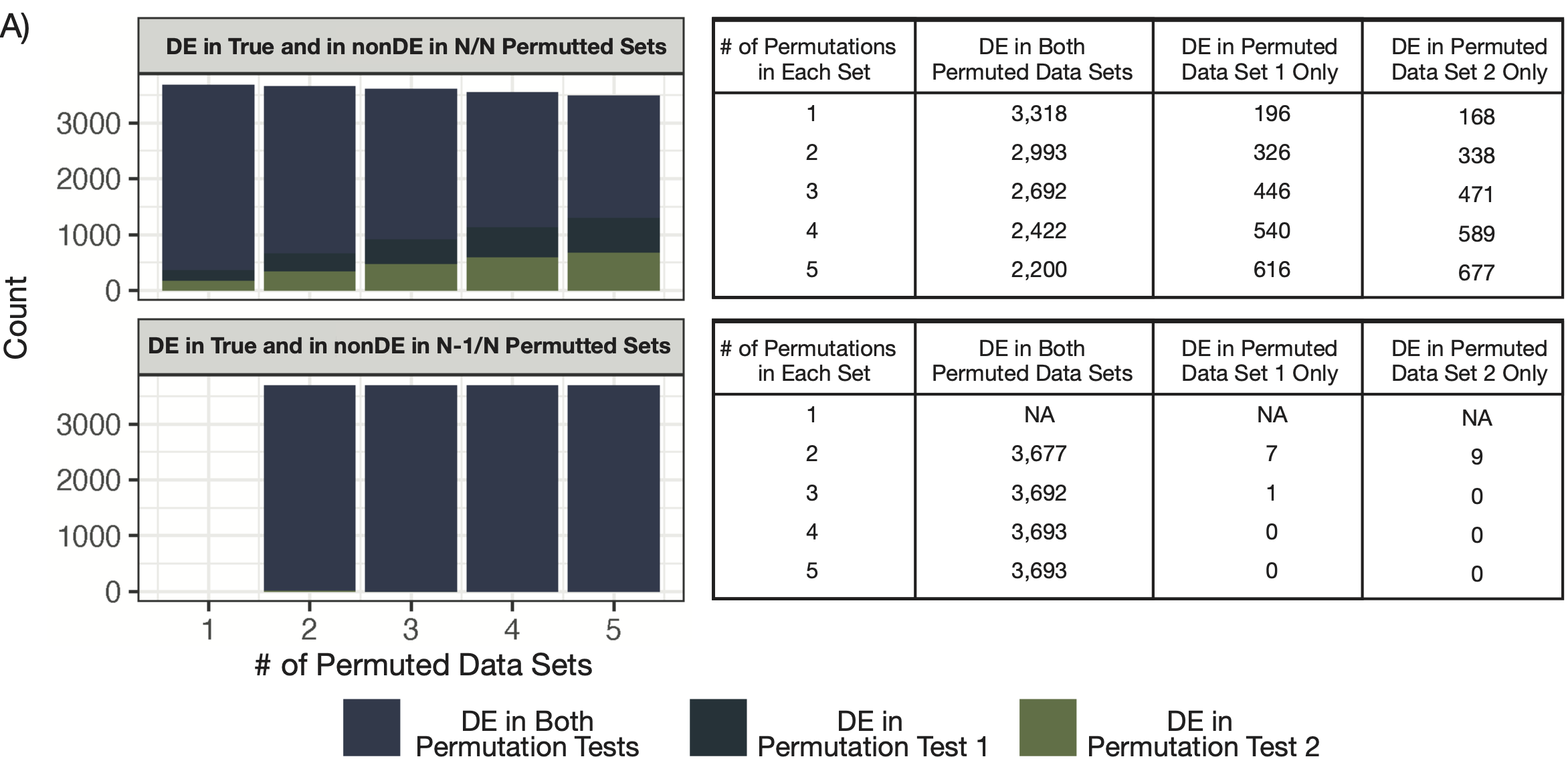

**Figure S8. Determining Appropriate DE Thresholds from Permutations**. We relied on a permutation-based approach to determine whether genes were differentially expressed in our true data set. We tested both the number of permutations (1 – 5) and how significance was assessed. Significant differentially expression was determined either by requiring the gene to be differentially expressed in the true data set and never in all the tested permuted data sets (i.e., non-differentially expressed in N/N permuted data sets) or by requiring the gene to be differentially expressed in the true data set and differentially expressed in at most 1 permuted data set (i.e., non-differentially expressed in at least N-1/N permuted data sets). We have two data sets each containing five permutations of our true data set. (A) A stacked barchart showing how many genes were called as differentially expressed in both of our tested data sets or in just one. The x-axis shows the number of permutations from the data set used, while the y-axis shows the number of genes. We faceted our results by how we assessed significance. Since numbers were sometimes very small, a table showing the numerical values was placed to the right of this figure.
